# Insitutype: likelihood-based cell typing for single cell spatial transcriptomics

**DOI:** 10.1101/2022.10.19.512902

**Authors:** Patrick Danaher, Edward Zhao, Zhi Yang, David Ross, Mark Gregory, Zach Reitz, Tae K. Kim, Sarah Baxter, Shaun Jackson, Shanshan He, Dave Henderson, Joseph M. Beechem

**Author notes:** **Corresponding author** Patrick Danaher, NanoString Technologies, Seattle, WA 98109, USA.

## Abstract

Accurate cell typing is fundamental to analysis of spatial single-cell transcriptomics, but legacy scRNA-seq algorithms can underperform in this new type of data. We have developed a cell typing algorithm, Insitutype, designed for statistical and computational efficiency in spatial transcriptomics data.

Insitutype is based on a likelihood model that weighs the evidence from every expression value, extracting all the information available in each cell’s expression profile. This likelihood model underlies a Bayes classifier for supervised cell typing, and an Expectation-Maximization algorithm for unsupervised and semi-supervised clustering. Insitutype also leverages alternative data types collected in spatial studies, such as cell images and spatial context, by using them to inform prior probabilities of cell type calls. We demonstrate rapid clustering of millions of cells and accurate fine-grained cell typing of kidney and non-small cell lung cancer samples.

## Introduction

A cornerstone of single cell transcriptomics analysis is cell typing, the process of categorizing cells based on their expression profiles. Cell typing can be performed using unsupervised clustering, which seeks to discover the cell types present in a sample. In tissues with cell types characterized by high-quality reference datasets (Regev 2017, Quake 2021, Allen Institute for Brain Science 2011), cell typing can employ supervised classification, avoiding the instability and interpretation difficulties of clustering. A hybrid approach, semi-supervised cell typing, detects known cell types while also discovering new clusters (Zanini 2020).

Spatial transcriptomics platforms detect individual RNA molecules where they lie on tissue slides, producing datasets of single-cell gene expression data while preserving cells’ location data (He 2022, Wang 2018). A typical experiment will measure 50-1000 genes and 10^5^ to 10^7^ cells (Anderson 2022). Cell typing is fundamental to analysis.

Existing approaches to cell typing spatial transcriptomics data fall into three classes. One class harnesses cells’ spatial locations to improve cell typing accuracy. Some of these methods model the frequency of each cell type within cells’ neighborhoods (Teng 2022), or within spatial domains (Chidester 2022, Li 2022). Others merge neighborhood information with the spatial data, appending neighborhood expression to the single-cell expression matrix prior to clustering (Singhal 2022) or by considering spatial information in the construction of a distance matrix between cells (Avesani 2022).

A second class of methods analyzes the locations of individual transcripts (Park 2021, Qian 2020, He 2021). These methods seek to simultaneously assign RNA molecules to cells and to assign cells to cell types. By forgoing input from increasingly powerful cell segmentation algorithms (Petukhov 2022, Greenwald 2022), these methods must fit more complicated models over single-molecule data, which is typically hundreds of times larger than single-cell data, posing a challenge for computation time.

A third class of methods uses spatial transcriptomics datasets as references for cell typing future datasets from the same platform (Brbic 2021). This class will become increasingly important as spatial data becomes more abundant.

Here we introduce the Insitutype algorithm for cell typing in spatial transcriptomics data. Insitutype is a built atop a likelihood model; this framework enables supervised cell typing from reference datasets via a Bayes classifier, and unsupervised and semi-supervised cell typing via an Expectation Maximization (EM) algorithm (Fig. 1). This ability to run with or without reference data makes Insitutype applicable to a wide range of studies.

**Figure 1.**
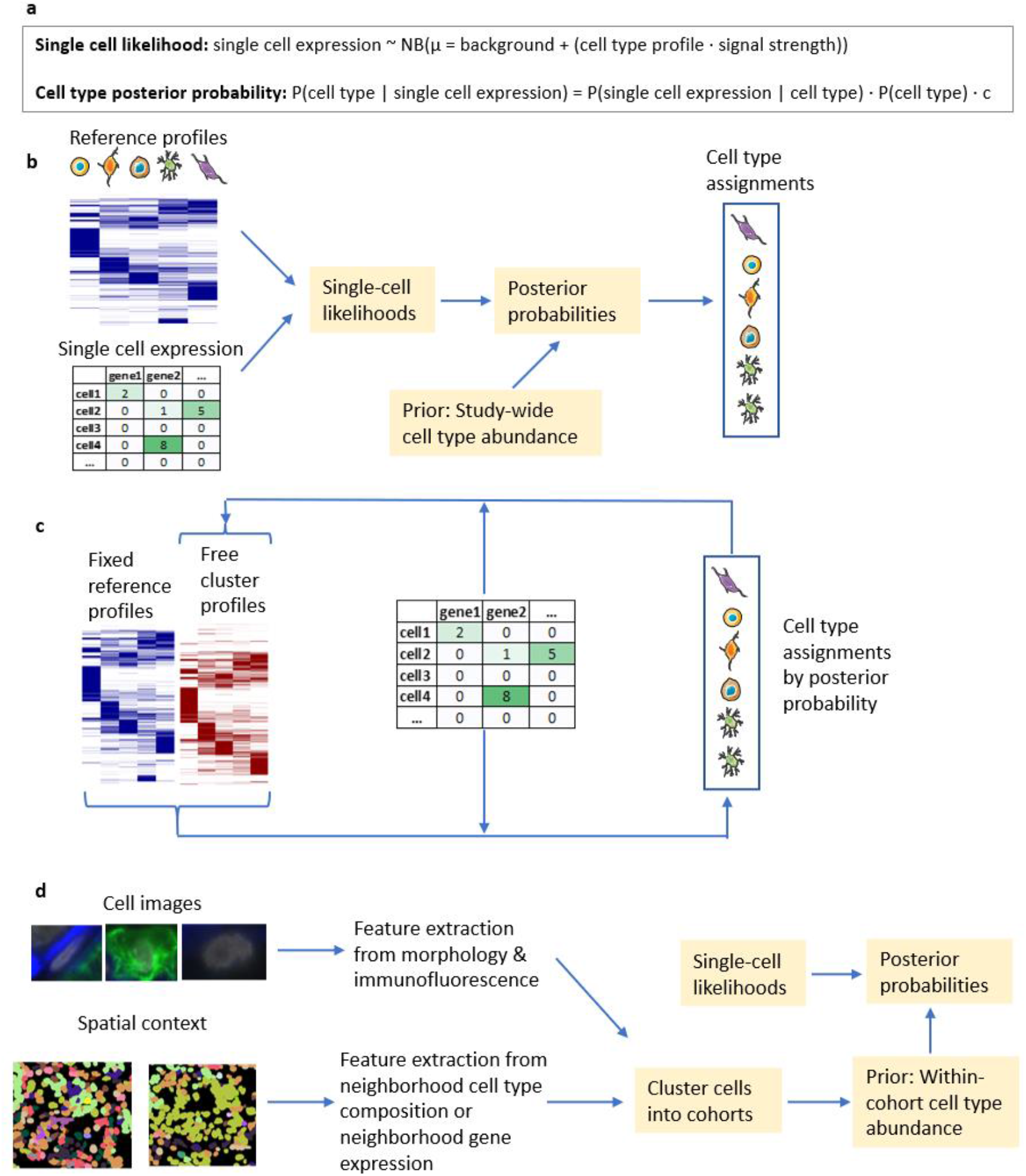
Insitutype overview. **a**. Single cell profiles are modeled with a negative binomial distribution whose mean depends on cell type, background and signal strength. Bayes’ rule gives the posterior probability of a cell type call. (c is a constant.) **b**. Supervised cell typing evaluates each cell’s likelihood under pre-specified “reference” cell type profiles, then uses these values to update prior cell type probabilities derived from study-wide cell type abundance. **c**. Unsupervised cell typing uses an EM algorithm to iterate between assigning cell types and estimating cell type profiles. Semi-supervised clustering holds reference profiles fixed while additional “free” profiles are updated by the EM algorithm. **e**. Alternative data types are used to cluster cells into cohorts, which then inform the likelihood calculation.

Insitutype is motivated by the challenges and opportunities of spatial transcriptomics data. First, compared to scRNA-seq, spatial transcriptomics produces sparser data: only a fraction of the transcriptome’s genes are measured, and RNA transcripts are missed in cells whose volumes extend beyond the thin tissue slide. With fewer transcripts per cell measured, it becomes important to use all available information with maximal statistical efficiency. Most scRNA-seq clustering methods employ dimension reduction, which reduces noise and improves computation time, but discards information. Popular approaches include partitioning a graph of nearest-neighbor distances (Traag 2019) or fitting cluster-specific centroids from the low-dimensional representation (Lloyd 1982, Scrucca 2016, Kiselev 2017). On the other hand, popular supervised scRNA-seq cell typing methods generally either rely on marker gene information (Zhang 2019, Cao 2020), which discard the information from most genes, or employ correlation-based metrics (Stuart 2019, Aran 2019, de Kanter 2019). Neither category models individual gene counts. In contrast, Insitutype’s likelihood framework is designed to optimally weigh the evidence from every transcript in a cell.

Second, spatial transcriptomics data is multi-modal. In addition to gene expression, it collects images of cells, and cellular neighborhoods act as a third data type. To exploit all available information, Insitutype is designed to incorporate alternative data types like image data and spatial context.

Finally, with spatial transcriptomics datasets of millions of cells becoming commonplace, computational efficiency is important. Insitutype employs an escalating subsampling scheme, attaining computation time that scales better-than-linearly and handles datasets of millions of cells.

Compared to existing cell typing methods, Insitutype has several advantages. It scales to large datasets that are intractable for some methods, it gains statistical power by modelling the likelihood of cells’ full expression profiles, it can identify new clusters alongside reference cell types, and it alone harnesses data from cell images. Thus far the cell images accompanying spatial transcriptomics data have only been used for imputation and spatial clustering, not as part of cell typing (Tang 2022).

Insitutype is available at https://github.com/Nanostring-Biostats/InSituType.

## Results

### A data-generating model of single-cell spatial transcriptomic data

The core of Insitutype is a likelihood model for spatial transcriptomics data. This model posits that for any given gene and single cell, the number of transcripts observed arises from a negative binomial distribution. The mean of this distribution depends on each cell’s sampling efficiency and background level, and on the expected expression within a given cell type.

We assessed Insitutype’s model in a cell pellet array profiled using the CosMx™ Spatial Molecular Imager. The observed count data agreed almost perfectly with the theoretical quantiles implied by the Insitutype model (Fig. 2a,b).

**Figure 2:**
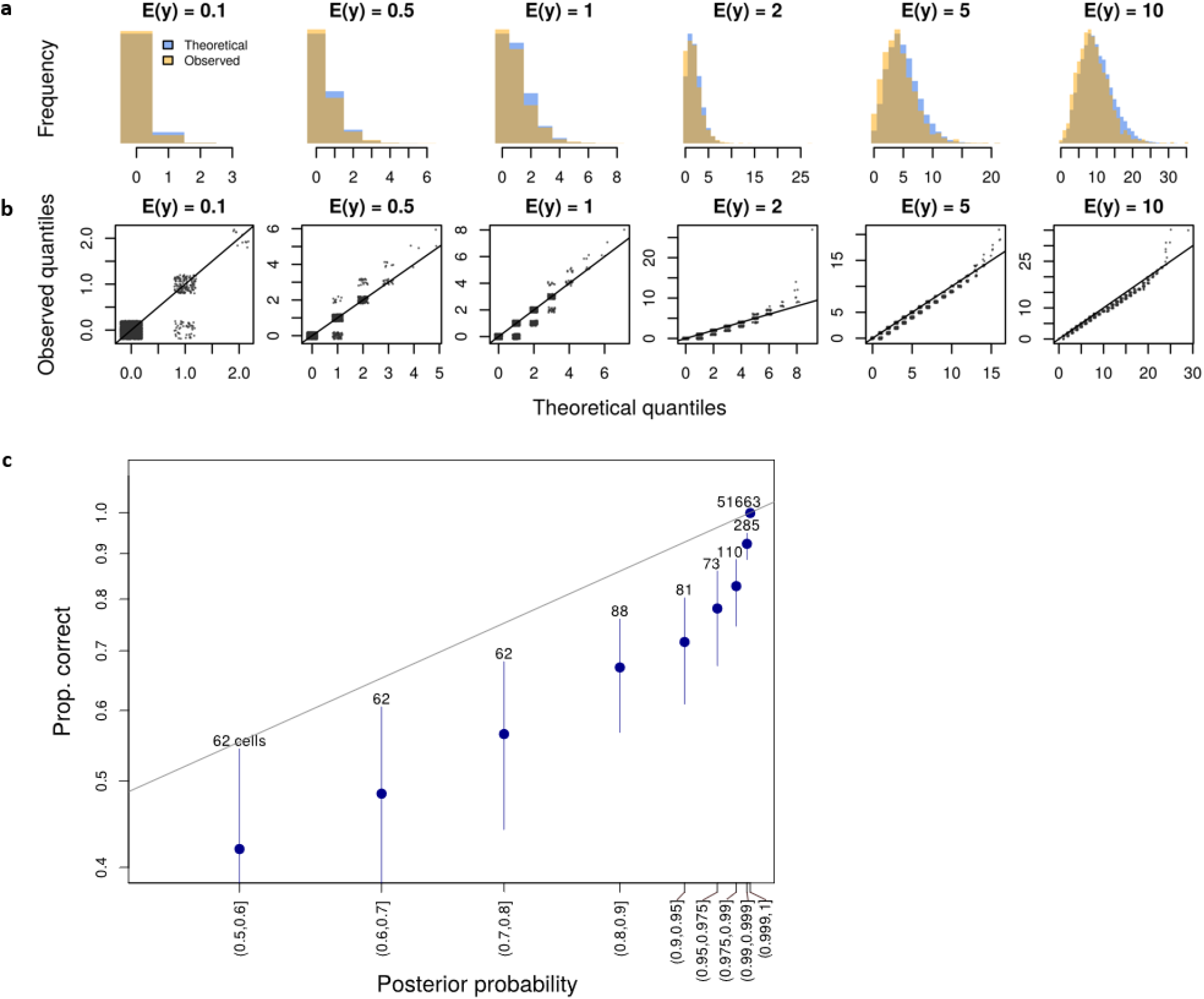
distribution of spatial transcriptomics data. For 6 expected expression levels, the 2000 data points (cells x genes) with expected counts closest to that level were selected. The observed counts at these data points were compared to the theoretical quantiles under Insitutype’s model. **a**. Distributions of expected and observed counts. **b**. Quantile-quantile plots of expected and observed counts. **c**. Accuracy stratified by posterior probability. Cells were divided into bins based on the posterior probability of their cell typing calls. Points show the rate of accurate cell typing calls within each bin; lines show 95% Wilson confidence intervals. Dashed line shows the identity. Numbers show number of cells within each bin of posterior probabilities.

### Quantifying cell typing uncertainty

Each Insitutype cell type assignment is accompanied by a posterior probability. These values can be used to remove low-confidence cell typing results from downstream analysis. To test the utility of posterior probabilities in flagging cell typing errors, we performed supervised cell typing within the cell pellet array shown in Fig. 2. Among the 85% of cells with posterior probability equal to 1, only 3 of 44,834 cell type calls were in error. As posterior probability decreased, so did accuracy (Fig. 2c). When cell typing calls were uncertain, accuracy was lower than the posterior probability suggested.

### Subsampling enables rapid analysis of millions of cells

To control computation time in large datasets, Insitutype’s EM algorithm uses an escalating subsampling scheme (Rocke 2003). To quickly approach an approximate solution, early iterations are run on small subsamples of 10,000 cells. Next, subsamples of 20,000 cells are used to obtain a more precise solution. The third set of iterations uses 100,000 cells, enough for a precise solution but far smaller than the number of cells in most datasets. The complete set of cells is analyzed only once, where they are classified using the profiles obtained from the 100,000-cell iterations. To enhance detection of rare cell types, Insitutype performs subsampling using geometric sketching (Hie 2019) (Supplemental Fig. 1).

Using this approach, Insitutype clustered 2,268,590 cells into 8 clusters in 30 minutes and 10 seconds running on a r5.12xlarge EC2 instance.

### Harnessing alternative data improves cell typing accuracy

When alternative data types such as cell images and spatial context are available, they can be used to partition cells into cohorts, either via clustering or by manual gating; for example, cells could be partitioned by CD45+/- status. Then, when analyzing gene expression data, Insitutype uses the prevalence of each cell type within each cohort to inform its posterior probability calculations (Methods).

To assess the impact of alternative data types, we clustered the data from the cell pellet array shown in Fig. 2, first using only gene expression data, second using cohorts derived from four immunofluorescence stains, and third using uninformative cohorts assigned at random. Cohorts can be defined flexibly and can benefit from careful attention. We simply fed the immunofluorescence data into Insitutype’s default cohorting function, which maps features onto Gaussian distributions then partitions cells into 100 clusters using model-based clustering. The average cell line’s error rate declined by a factor of 0.69 when the immunofluorescence data was incorporated, from 0.447% (Wilson method 95% confidence interval: 0.394%-0.508%) to 0.281% (0.240%-0.331%). Use of uninformative cohorts left error rate effectively unchanged, at 0.440% (0.387%-0.500%) (Fig. 3a). The impact was greatest in cells with fewer transcripts. Similar results hold for supervised cell typing, where the average cell line’s error rate declined by a factor of 0.65 when immunofluorescence data was used, from 0.508% (0.394%-0.508%) to 0.331% (0.286%-0.384%) (Fig. 3b).

**Figure 3.**
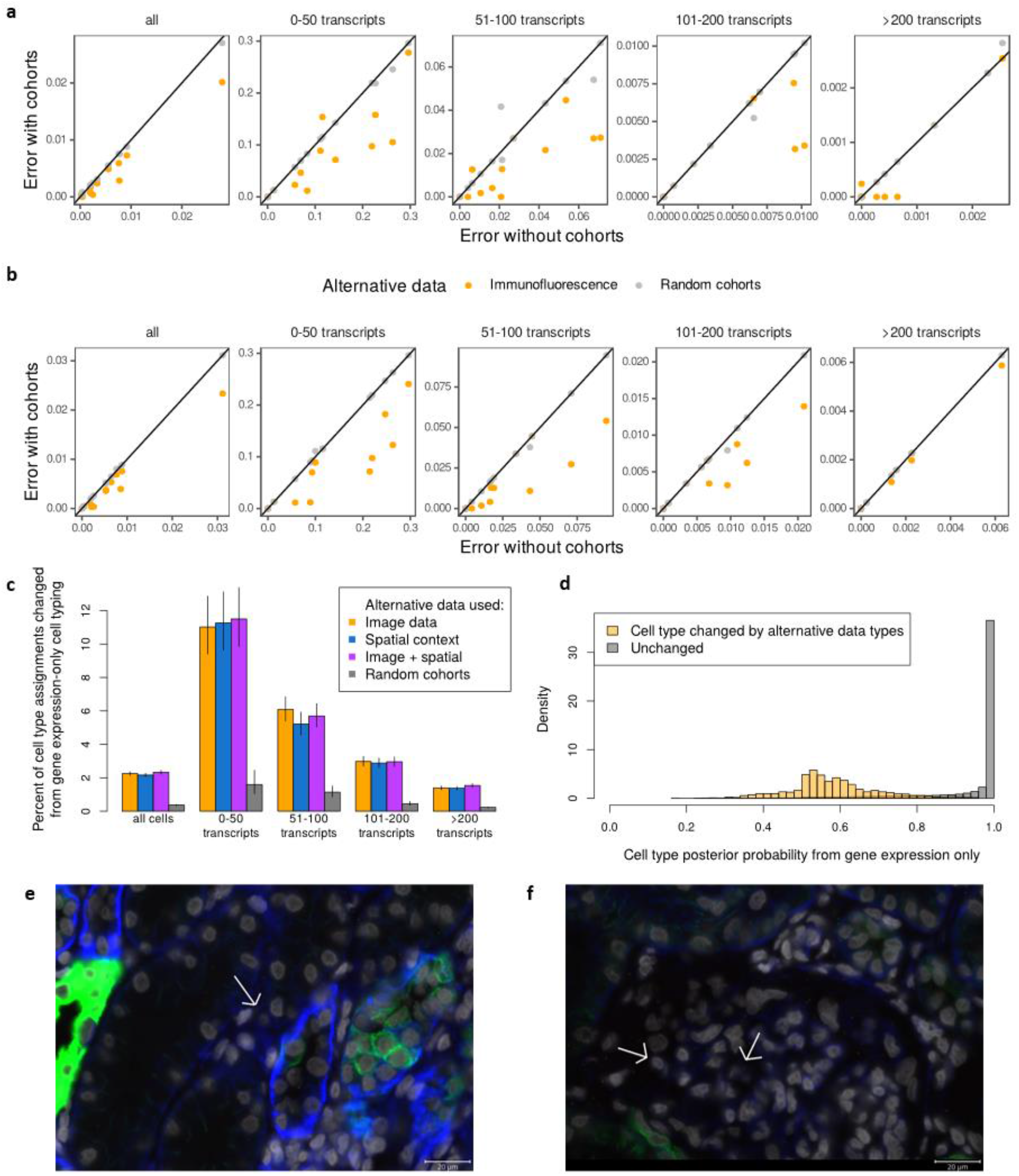
Impact of alternative data types. **a**. In unsupervised clustering of data from 13 cell lines, error rates when omitting alternative data (x-axis) and when incorporating alternative data as cohorts (y-axis). **b**. In supervised cell typing of data from 13 cell lines, error rates when omitting alternative data (x-axis) and when incorporating alternative data as cohorts (y-axis). **c**. Impact of alternative data types on supervised cell typing in a kidney sample. Bars show percent of cells; lines show 95% confidence intervals. **d**. Strength of RNA evidence for cell type assignments changed/unchanged by immunofluorescence and spatial data. Histograms show density of RNA-only cell typing posterior probabilities across all cells. **e**. A cell called a podocyte by RNA only but called a fibroblast when image and spatial data were used. **f**. Two cells called “ascending vasa recta endothelium” by RNA only and called “glomerular endothelium” when image and spatial data were used. Both cells are in a glomerulus.

We then investigated the impact of alternative data on cell typing in a lupus nephritis sample. First, image data was extracted for each cell, including area, aspect ratio and mean intensity of five immunofluorescence markers. Second, cells’ spatial context was defined using the mean expression of all neighboring cells within a 50µm radius (Methods). Cell typing was performed with RNA alone, with RNA and images, with RNA and spatial data, and with all three data types. Image data and spatial context each changed 2.2% of cells’ assignments (Fig. 3c), while random cohorts changed only 0.4% of assignments. The alternative data types mainly impacted cells with limited gene expression data: among cells that changed assignment with the use of image data and spatial context, only 4% had posterior probability from gene expression >0.8; among cells that did not change assignment, 90% had posterior probability >0.8. (Fig. 3d). When image and spatial data overrode an RNA-only cell type assignment, it was often in clear agreement with known biology, for example correcting a “podocyte” label outside a glomerulus (Fig. 3e), or reclassifying cells inside a glomerulus from “ascending vasa recta endothelium” to “glomerular endothelium” (Fig. 3f).

### Benchmarking Insitutype against other cell typing algorithms

We benchmark the accuracy of Insitutype against other unsupervised clustering algorithms (Fig. 4a-c) and supervised cell typing algorithms (Fig. 4d-f) using a cell pellet array dataset of 52,518 cells. Benchmarking is performed on the full dataset, as well as on two downsampled datasets containing a random sample of 50% of the genes and the 10% most highly variable genes.

**Figure 4.**
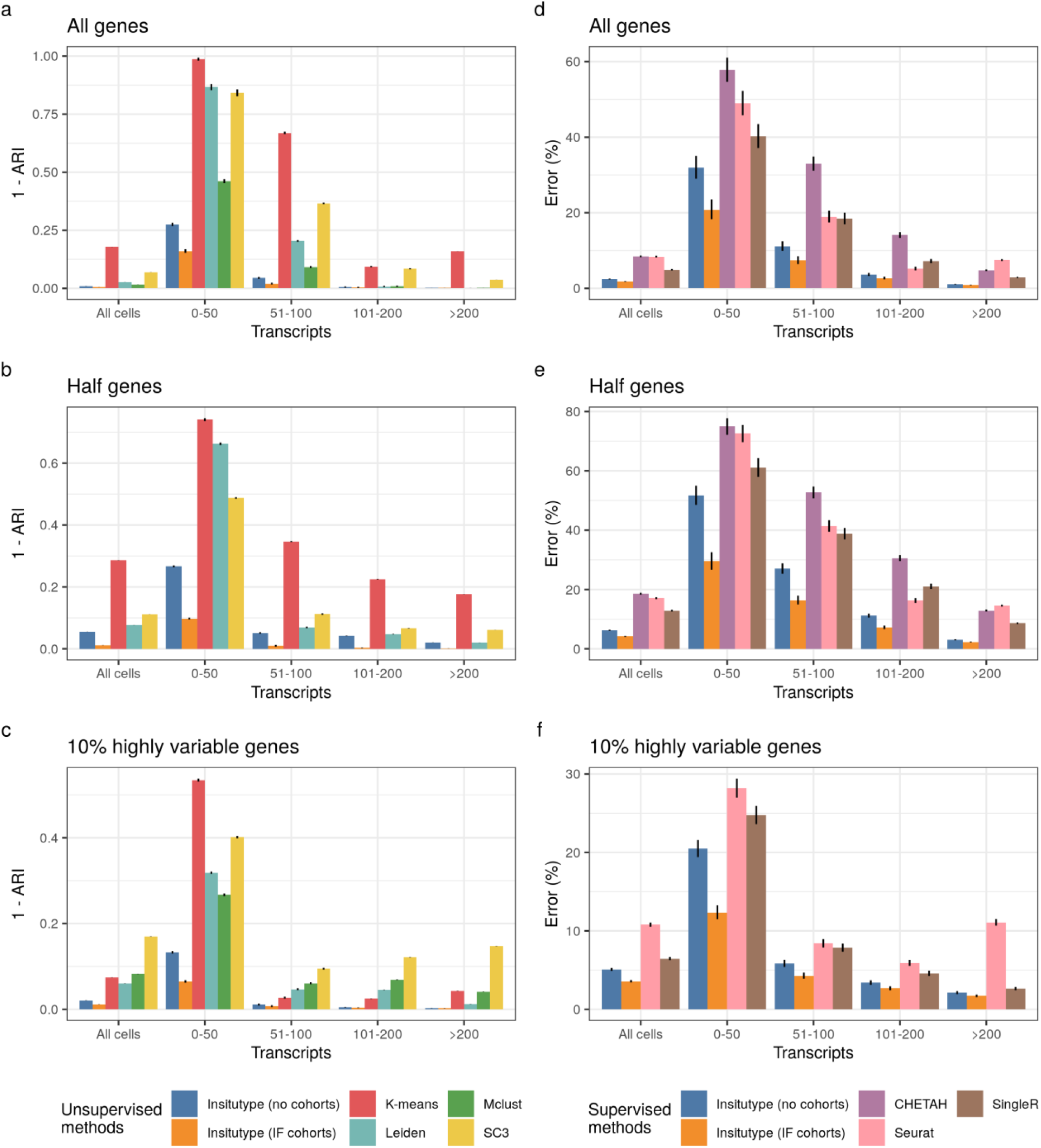
Benchmarking performance. **a**. Unsupervised clustering of data from 13 cell lines using all genes, **b**. using half of the genes randomly sampled, and **c**. using the top 10% most highly variable genes. Bars show 1 – adjusted Rand index (ARI) to compare the inferred clustering partition with the true cell line. Lines show 95% confidence intervals. Groups of bars summarize cells with similar total transcript counts. **d**. Supervised cell typing of data from 13 cell lines using all genes, **e**. using half of the genes randomly sampled, and **f**. using the top 10% most highly variable genes. Bars show the error rate and lines show 95% Wilson confidence intervals. Groups of bars summarize cells with similar total transcript counts.

Among unsupervised methods, Insitutype consistently outperforms K-means, Mclust, Leiden, and SC3. The use of immunofluorescence cohorts further improves Insitutype’s performance, and this effect is more noticeable for cells that contain fewer transcripts. Downsampling genes impacted these methods differently. For example, Insitutype performance is comparable across the three choices of gene selection, but k-means performance is substantially better when using highly variable genes (Fig. 4a-c). Mclust failed to converge using principal components derived from 480 randomly selected genes.

For supervised cell typing, we generated a reference dataset by assaying the cell pellet array on a different instrument. Insitutype makes consistently fewer errors than CHETAH, Seurat, and SingleR. The addition of immunofluorescence data again improves Insitutype’s accuracy. CHETAH failed to run when using only highly variable genes.

### Supervised cell typing of a lupus nephritis biopsy

To assess Insitutype’s ability to perform supervised cell-typing in real tissues, 960-plex CosMx data was collected from a kidney core biopsy from a lupus nephritis patient. Insitutype assigned the biopsy’s 61,073 cells to 33 cell types based on gene expression, cell images and neighborhood context (Fig. 5a).

**Figure 5:**
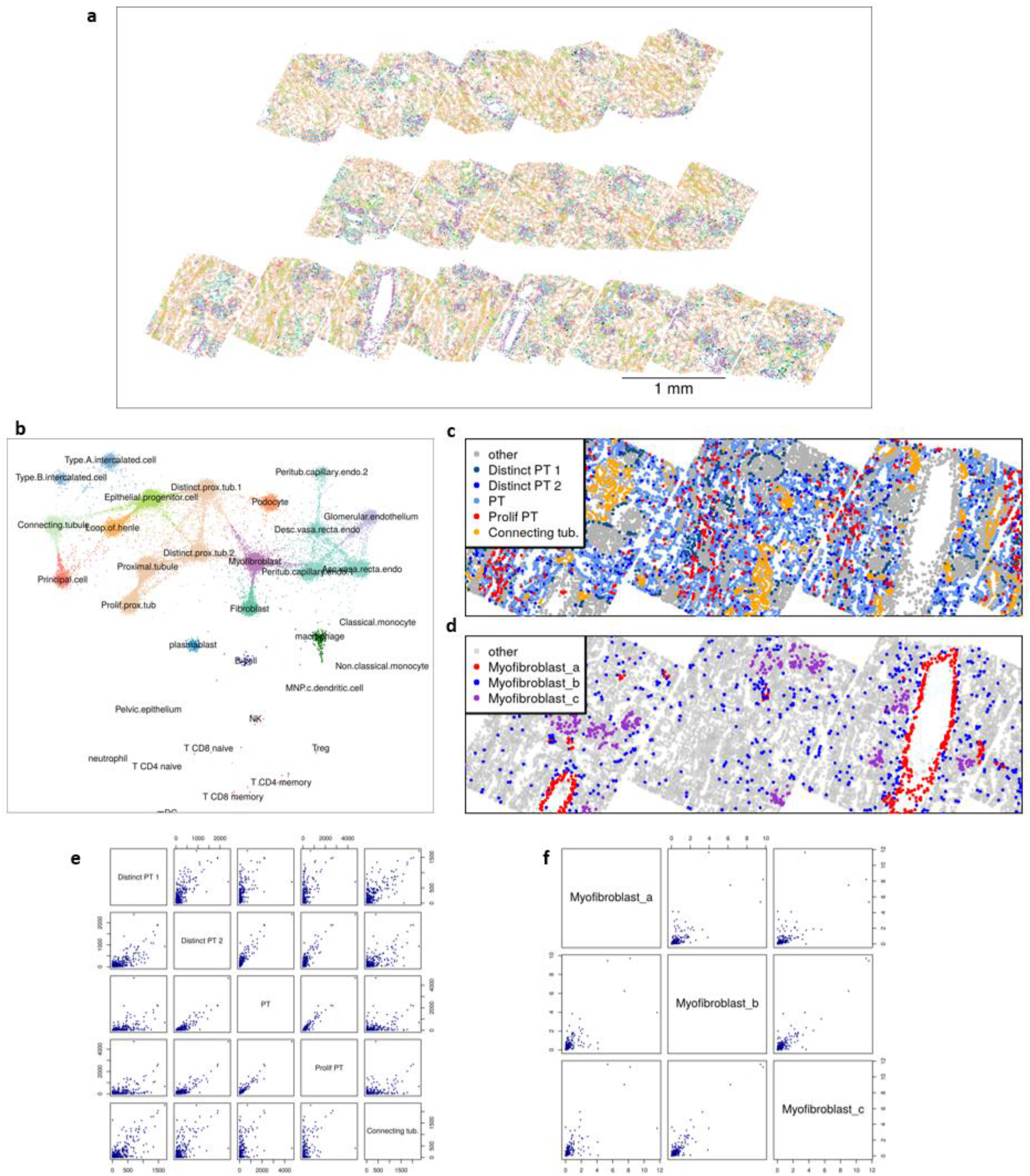
Supervised cell typing in a lupus nephritis kidney biopsy. **a**. Map of cell types across the tissue. Cell type colors are defined in (b). **b**. Cell typing confidence. Points are positioned according to their posterior probability of belonging to each cell type. **c**. Proximal tubule subtype calls. **d**. Myofibroblast sub-cluster calls. **e**. Reference profiles of proximal tubule subtypes. **f**. Profiles of myofibroblast sub-clusters.

Cell typing confidence in summarized in Fig. 5b. This view, which we term the “flightpath” plot, places cells according to their posterior probabilities of belonging to each cell type. This view conveys the tendency of different cell types to be confused with each other; we propose it as a useful summary of any cell typing algorithm’s results. In this tissue, most cell types were discerned from each other with high confidence, while higher confusion rates were seen among closely related cell types, for example the proximal tubule subtypes and the endothelial cell subtypes.

Spatial patterns help us assess the ability of Insitutype to discern closely related cell subpopulations. The Human Cell Atlas reported four subtypes of proximal tubules, the dominant cell type in kidneys. Insitutype finds these subtypes to have distinct spatial distributions (Fig. 5c,e). “Proximal tubule” cells dominate, and “distinct proximal tubule 2” cells are scattered among them at lower frequency, while “distinct proximal tubule 1” cells surround glomeruli. Based on their spatial distribution, this last cluster can now be identified as parietal epithelium cells. “Proliferating proximal tubule” cells fall in bands consistent with ducts injured by autoimmune attack. Meanwhile, “connecting tubule” cells lie in clusters and bands consistent with tubular morphology.

The Human Cell Atlas reported a cell type, “myofibroblast”, that we find to encompass both typical myofibroblasts and specialized myofibroblasts found only in glomeruli, called mesangial cells. Insitutype was used to sub-cluster the study’s 6768 myofibroblasts. The resulting 3 clusters recovered mesangial cells concentrating in glomeruli, interstitial myofibroblasts scattering diffusely, and vascular pericytes ringing vasculature (Fig. 5d,f).

### Semi-supervised cell typing of non-small cell lung cancer

The NanoString CosMx FFPE non-small cell lung cancer public data release reports 960-plex RNA data from 771,236 cells across 5 tumors. We applied Insitutype’s semi-supervised approach to this dataset, calling previously defined immune and stroma cell types (Danaher 2022) while detecting additional cell types missing from the reference profiles.

From the above inputs, Insitutype called cells from all 16 reference cell types and from 10 additional clusters. The unsupervised clusters included 6 tissue-specific clusters of cancer cells, a healthy epithelium cluster shared across tumors, a second macrophage cluster with enriched DUSP5 and lower SPP1, a second plasmablast cluster with lower IgG gene expression, a second fibroblast cluster with enriched DUSP5 and reduced MMP1, and a non-specific cluster drawing from multiple stroma cell types (Fig. 6a,b, Supplemental Fig. 3,4,5). Cells initially assigned to the non-specific stroma cluster were reassigned to the cell type providing the next best fit. Cell typing confidence was high for most cells: excluding the T-cell subclusters, NK cells, pDCs and monocytes, the remaining cell types had >0.9 posterior probability for over 80% of their cells.

**Figure 6:**
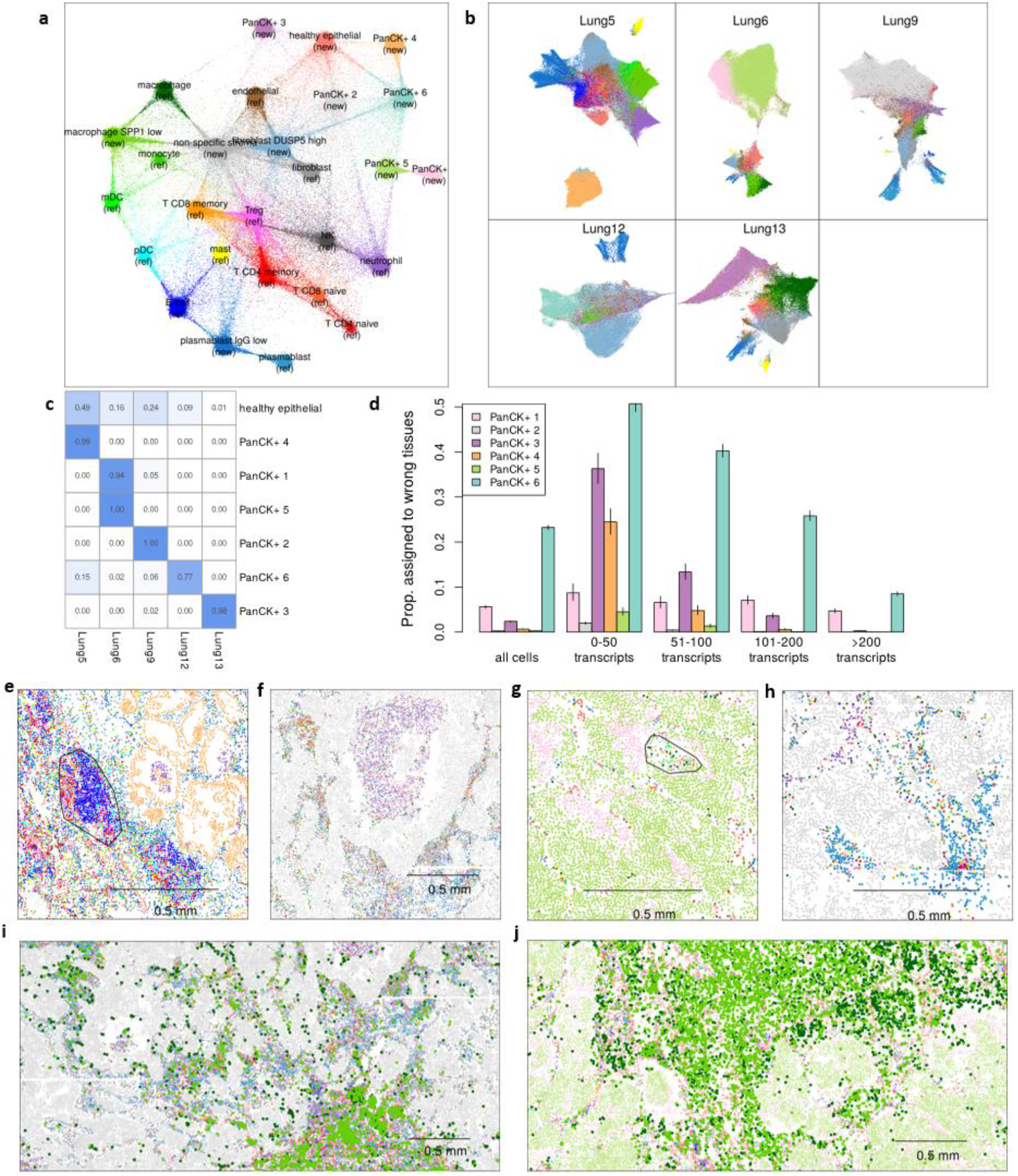
Semi-supervised cell typing in non-small cell lung cancer. **a**. Cell typing confidence. Points are positioned according to their posterior probability of belonging to each cell type. Colors are consistent with panels b and d-j. **b**. UMAP projections of each sample, colored by cell type. **c**. Frequency of each tissue within each PanCK+ cluster. **d**. Error rates of each tumor-specific PanCK+ cluster. Lines show Wilson method 95% confidence intervals. **e**. A putative tertiary lymphoid structure. **f**. a pocket of neutrophils. **g**. macrophages and T-cells invading the tumor bed. **h**. Plasmablasts spreading out from a core of T-cells. **i**. SPP1-high and low macrophages in Lung9. **j**. SPP1-high and low macrophages in Lung6.

To assess clustering accuracy, we examined the PanCK+ clusters, which each appeared to represent a single tumor cluster from a single patient. When a tumor-specific cluster includes cells from another tissue, it is fair to call these assignments errors. Most tumor-specific clusters contained >94% of cells from the correct tissue (an error rate of 6%), although one cluster had only 77% of cells from the correct tissue (Fig. 6c). When only cells with >200 transcripts are considered, error rates decrease to <0.27% for most clusters and 8.5% for the worst-performing cluster (Fig. 6d).

The cell type map returned by Insitutype agrees with and expands on the known behavior of immune cells in lung cancer. Complex immunologic phenomena can be seen, for example tertiary lymphoid structures, pockets of dense neutrophils, T-cells and macrophages invading deep into the tumor, and large gatherings of plasmablasts alongside a small core of T-cells (Fig. 6e-h).

Two macrophage sub-populations were identified (Supplemental Fig. 3). The macrophage reference profile identified macrophages with elevated SPP1, a gene linked to immunosuppressive tumor-associated macrophages (TAMs) (Zhang 2017). A de novo cluster, presumably classical macrophages, was SPP1-low. These subpopulations concentrated differently within and across tissues. When both populations were prevalent in a tissue, they displayed distinct spatial distributions. In Lung9, SPP1-high macrophages were primarily found in the tumor bed, while SPP1-low macrophages gathered densely in the stroma (Fig. 6i). In Lung6, both populations were found in the stroma, but they aggregated in distinct areas within the stroma (Fig. 6j). Thus in five samples we see four distinct patterns of tumor-associated and classical macrophage distribution: TAMs were all but absent in two samples, dominant in another sample, predominantly confined to the tumor bed in one sample, and abundant in both tumor and stroma regions of the final sample.

### Comparison to other spatial clustering methods

We used a NSCLC sample from Figure 6 (Lung5) to compare Insitutype to alternatives. Of the published clustering methods designed for spatial transcriptomics, we were only able to run BASS on this dataset. BASS derives spatial domains alongside its single cell type assignments. We used both BASS and unsupervised Insitutype to fit 12 clusters to the sample’s 97,809 cells (Supplemental Fig. 6). The two methods agreed on most cell types, but diverged in where they fit finer sub-clusters. Insitutype partitioned the tumor cells into 3 clusters, while BASS found more granular partitions of the immune cells. In this relatively small dataset, Insitutype ran in 8 minutes; BASS took 7 hours and 10 minutes.

## Discussion

We propose Insitutype as a cell typing solution applicable to any single cell spatial transcriptomics study, regardless of plexity, platform (Supplemental Fig. 2), number of cells, or availability of reference datasets. Insitutype’s likelihood framework and ability to incorporate alternative data types like cell images and spatial context enhance statistical power; we have shown how this power enables fine-grained cell typing in clinical FFPE specimens.

Conceptual innovations in this work include the use of a likelihood model to weight the evidence from every expression value, the explicit modeling of background, the use of cohorts to incorporate alternative data types with diverse distributions, the use of subsampling to speed algorithm convergence in large single cell datasets, the use of single cell images to inform cell typing, and the useful display of clustering results by posterior probability shown in Figures 5b and 6a. Biological findings include the discovery of cell types missing from the kidney scRNA-seq reference, and the presence of the IgG-low plasmablast subtype and the DUSP5-high fibroblast subtype across multiple tissues.

Insitutype’s semi-supervised capability can be indispensable in spatial transcriptomic analyses. scRNA-seq reference datasets often miss hard-to-dissociate cell types, leaving additional clusters to be discovered in spatial studies. And in tumors, while immune reference profiles can be pre-defined, the high inter-patient heterogeneity of cancer cells defies use of reference profiles.

Insitutype uses alternative data types to define “cohorts” of cells; cohort membership then defines cells’ prior probability of belonging to each cell type, and thereby influences their cell type posterior probabilities. By incorporating alternative data types in this manner, Insitutype appropriately weighs their evidence without reliance on ad hoc tuning parameters. This cohorting approach is convenient: whatever alternative data is available, cohorts are defined according to the best judgement of the analyst, and these cohorts’ impact on cell typing increases with the information they hold. However, by condensing alternative data into categorical cohorts rather than modelling its continuous values, Insitutype fails to fully exploit the information in the alternative data. Insitutype’s default approach mitigates this concern by defining hundreds or thousands of fine-grained cohorts, capturing the information from alternative data types as thoroughly as possible with a categorical variable.

Spatial context offers useful clues about cell type identity, which Insitutype exploits through cohorting. However, if the intent is to study how gene expression or cell type abundance changes across spatial contexts, then using spatial context in cell typing risks introducing artifacts into the analysis. It will often be better to accept a slightly higher error rate and avoid concerns about circular reasoning.

A limitation is that Insitutype models each cell type with a single centroid, ignoring the possibility of varying expression states within a cell type. Future extensions could model each cell type with a manifold instead of a centroid. In a close approximate of this strategy, Insitutype can be run with high cluster numbers; closely-related clusters can then be merged into a single cell type.

In Figures 5b and 6a, we plot cells according to their posterior probabilities of belonging to each cell type. We have found this view, which we have named the “flightpath” layout, to be indispensable. Although this plot depends on log-likelihoods output by Insitutype, it can be applied to results of other clustering methods. Given cluster assignments from an arbitrary method, the required log-likelihood matrix can be derived by first calculating the clusters’ average background-subtracted expression profiles, and then feeding these profiles into Insitutype’s supervised cell typing workflow.

In our kidney analysis, we discovered cell types missing from the Human Cell Atlas reference: the “myofibroblast” cell type held three distinct sub-clusters, and the “distinct proximal tubule 1” cell type contained parietal epithelial cells. The ability of spatial transcriptomics to discover cell types missed by scRNA-seq has two explanations. First, some cell types could have failed to dissociate from tissue and so were missed entirely by scRNA-seq. This explains the three myofibroblast subtypes, which had highly distinct expression profiles. Alternatively, cell types with very similar transcriptional profiles can be more easily discerned if spatial context is known. This finding suggests that cell atlases built on dissociated cells alone may be incomplete, and that spatial technologies can discover additional cell types.

## Acknowledgements

We are grateful to Martin Hemberg and Jingyi Cao for helpful feedback on the manuscript and software.

## Methods

### Notation

Let Y be the matrix of observed expression in *M* genes over *N* cells.

Let X be the expected expression profile of *M* genes across *K* cell types.

Let *i* index gene, *j* index cell, and *k* index the *K* cell types. Call *k(j)* the cell type of cell j.

### Likelihood model for gene expression in spatial transcriptomics

We model a cell’s true, unobserved gene expression as derived from a negative binomial distribution, with mean determined by cell type:

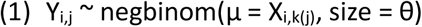

When we measure this expression with a spatial transcriptomics platform, it becomes distorted by three technical effects. First, not every mRNA transcript in the cell will be detected. We model the detection efficiency within a cell *j* as a scaling factor *s*_*j*_. These scaling factors will vary between cells as a function of both mRNA reading efficiency in the area containing the cell and the proportion of the cell’s volume present on the slide (cutting a slide from a tissue block will slice many cells in half).

Second, due to off-target binding, spatial transcriptomics platforms produce “background” counts, adding spurious counts to a cell’s observed expression profile. Call a cell’s expected number of background counts per gene *b*_*j*_.

Third, a given platform might measure different genes with different efficiencies. These gene-specific efficiencies can be incorporated into the model by rescaling the rows of X, the matrix of expected expression per cell type; therefore we do not add a term to the model capturing these factors.

Accounting for the above technical effects suggests the following model for observed gene expression:

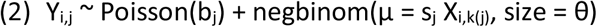

Modelling convolutions of discrete probability distributions can be computationally expensive. Therefore Insitutype assumes a simplified model:

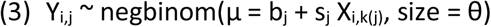

This simplification produces the same mean model as (2), and slightly higher variances for the same value of θ.

### Estimating individual cell’s background rates

Spatial transcriptomics platforms include negative control “genes” designed to assess background rates. Some platforms (He 2022) use physical negative controls - hybridization probes targeting sequences not present in any known genome (External RNA controls consortium 2005). Other platforms (Wang 2018) use computational negative controls, directing their software to detect barcodes not generated by any gene.

Within a given cell, the number of negative control counts will be zero or near zero. Therefore, due to the distribution of Poisson random variables with near-zero rates, negative controls are a noisy measurement of a cell’s background rate. For example, if a cell has a background rate of 0.04 counts per gene, then 10 negative control probes will usually return either 0, 1 or 2 total counts, corresponding to background rates of 0, 0.1 or 0.2, either suggesting that there is no background or estimating a background rate more than double the truth.

To attain a more stable estimate of cells’ background, Insitutype’s default is to predict a cell’s background from its total counts. It fits an intercept-free linear model predicting cells’ mean negative control gene expression from their total counts, and it treats this model’s fitted values as each cell’s background tendency *b*_*j*_.

### Estimating individual cell’s scaling factors

Given a cell’s observed expression Y_,j_, its expected background b_j_ and its expected expression vector X_,k(j)_, it is possible to compute a maximum likelihood estimate for s_j_ through diverse optimization algorithms. To avert the need to perform possibly millions of iterative optimizations, Insitutype approximates *s*_*j*_ as follows:

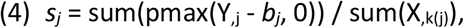

where pmax is the parallel maximum function, returning a vector over the genes, containing the larger of Y_i,j_ – b_j_ or 0 for gene *i*.

### Computing a cell’s log-likelihood under each possible cell type

The algorithm for classifying a cell based on its expression profile Y_,j_ and the cell type profile matrix X is as follows:

#### Algorithm 1

Assume *b*_*j*_ and θ have already been calculated.

For each cell type *k*:

1. Calculate *s*_*j*_ as described by Equation (4).
2. Compute the cell’s expected counts under cell type k as 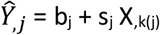
3. Calculate the cell’s log-likelihood as dnbinom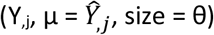, where dnbinom is the negative binomial log-likelihood function of equation (3).

### Updating cell log-likelihoods based on frequencies

For a cell *j*, Algorithm 1 gives us the likelihood of the profile Y_,j_ under cell type k(j). We use k to denote k(j) for simplicity:

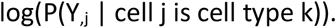

But we really desire to know the probability that the cell is cell type k given its profile:

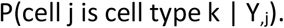

By Bayes’ theorem, we know:

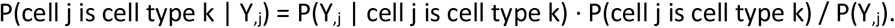

Let p_k_ be the proportion of all cells in the study that are cell type k, and use p_k_ to estimate the marginal probability that a cell is cell type k. Then we can write

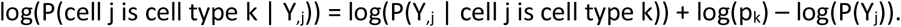

On the right side of the equation, the first term is output by Algorithm 1. The final term applies equally to all cell types *k*, and so can be ignored. Thus we calculate the posterior log-likelihood of a cell *j* belonging to a cell type *k* by adding log(p_k_) as follows:

#### Algorithm 2

Take as input the matrix of individual cells’ log-likelihoods under each cell type, output by Algorithm 1. Call this matrix “logliks”.

1. Classify each cell as whatever cell type gives it the greatest log-likelihood.
2. For all cell types:
  a. Compute “p_k_”, the frequency of cell type *k* in the output of step 2.
  b. Add log(p_k_) to column k of logliks.
3. The resulting matrix is the posterior log-likelihood of each cell under each cell type (with each row offset by a constant).

This use of a posterior log-likelihood instead of a conditional log-likelihood makes Insitutype more likely to assign cells to common cell types than to rare cell types. This means that rare cell types are only called with strong evidence, that analyses of rare cell types are not contaminated by numerous mis-classified cells of other types.

### Merging alternative data types into the likelihood model

This posterior probability calculation of Algorithm 2 can be harnessed to incorporate evidence from alternative data types like images and spatial context. Here we derive the procedure for doing so:

Notation:

- Let W_j_ be cell j’s data from alternative data types, e.g. a vector of features derived from the cell’s image and/or neighborhood details.
- Let “m_jk_” be the proposition that “cell j has cell type k”.

We desire P(m_jk_ | W_j_, Y_,j_), which equals P(m_jk_, W_j_, Y_,j_) / P(W_j_, Y_,j_).

Now P(m_jk_, W_j_, Y_,j_) = P(Y_,j_, m_jk_ | W_j_) P(W_j_).

And P(Y_,j_, m_jk_ | W_j_) = P(Y_,j_ | m_jk,_ W_j_) P(m_jk_ | W_j_).

And if we assume that a cell’s expression profile depends only on its cell type and not alternative data types of W, then we can claim P(Y_,j_ | m_jk,_ W_j_) = P(Y_,j_ | m_jk,_).

Note that this assumption will not hold perfectly for some cases. For example, if a cell changes expression profile based on its spatial context, and W includes spatial information, then the assumption will fail. However, even in this case, it is reasonable to assume that spatial context only slightly modifies the cell’s expression, i.e. that spatial context influences expression profile much less than cell type does.

Now, making all the above substitutions, we get

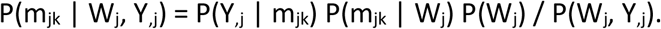

To assign a cell type, we only need to compare this expression across different cell types k. Thus terms that do not include m_k_ can be ignored:

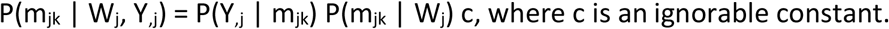

Then our posterior log-likelihood is

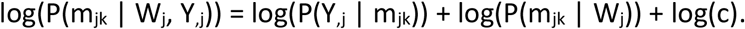

Since log(P(Y_,j_ | m_jk_)) is the output of Algorithm 1, this leaves only P(m_jk_ | W_j_) to be calculated in order to estimate log(P(m_jk_ | W_j_, Y_,j_)).

Since the data in W_j_ can be from diverse sources without obvious probability distributions, for example spatial context and cell images, a non-parametric estimate of P(m_jk_ | W_j_) is appropriate. To this end, Insitutype collapses all the information in alternative data types into a vector of “cohorts”, where each cohort contains cells with similar W data.

Cohorts may be defined in diverse ways. In its simplest version, the cohort vector could simply split cells based on positive / negative stains for a major immunofluorescence marker like Pan-cytokeratin. In a more elaborate treatment of image data, cell images could be transformed by an autoencoder into a low-dimensional matrix, and this matrix could be clustered into as many cohorts as desired. Spatial information can be encoded in diverse ways. Cells could be binned based on their x,y positions across a tissue. Or more informatively, a spatial clustering algorithm (Zhu 2018, Chidester 2022, Li 2022, Singhal 2022) could be used to cluster cells based on the gene expression profile in a small region around them, linking cells from the same spatial context wherever they fall within the tissue. To facilitate derivation of the cohort vector, the Insitutype library provides a function for automatically dividing cells into cohorts based on their immunofluorescence intensities and the gene expression profile of their spatial neighborhood.

Once the alternative data W has been collapsed into a vector of cohorts, calculating P(m_jk_ | W_j_) is trivial: its value is simply the frequency of cell type k within cell j’s cohort. To guard against sampling noise in small cohorts, when estimating P(m_jk_ | W_j_) we include 100 cells’ worth of abundances from the study-wide frequencies. For example if 1.5% of cells in the study are T-cells, then 1.5 T-cell counts will be added to each cohort’s abundances prior to calculating P(m_jk_ | W_j_).

### Estimating cell type expression profiles given cell type assignments

Insitutype uses the below algorithm to estimate the expression profiles of all cell types.

#### Algorithm 3

For each cell type:

1. Calculate mean signal as the vector of all genes’ mean expression across cells currently assigned to the cell type.
2. Calculate mean background as the mean of the matrix of negative controls across the same cells.
3. Subtract the mean background value from the mean signal vector, and threshold below at 0.

### Supervised cell typing

#### Algorithm 4

Supervised cell typing based on pre-specified reference profiles proceeds as follows:

1. Take as input:
  a. The observed expression matrix Y
  b. A pre-specified matrix of expected cell type expression profiles X
  c. Estimated per-cell background, likely estimated as described in the section, “Estimating individual cell’s background rates”.
  d. Mean negative control counts per cell.
  e. A vector specifying cells’ “cohort” membership, potentially all the same.
2. Compute conditional log-likelihoods using Algorithm 1.
3. For each cohort of cells, compute posterior log-likelihoods using Algorithm 2.
4. Assign each cell to the cell type under which it has the highest posterior log-likelihood.

### EM algorithm for unsupervised and semi-supervised clustering

Our basic clustering approach is an Expectation Maximization (EM) algorithm, described below. The full Insitutype algorithm orchestrates multiple calls to this basic EM algorithm.

#### Algorithm 5

Take as input:

a. The observed expression matrix Y
b. An initial matrix of expected cell type expression profiles X^(free)^ (potentially carefully pre-specified, or potentially derived from random cluster assignments.)
c. Estimated per-cell background, likely estimated as described in the section, “Estimating individual cell’s background rates”.
d. Mean negative control counts per cell.
e. Optionally, a matrix of fixed reference profiles X^(ref)^.
f. A vector specifying cells’ “cohort” membership, potentially all the same.

2. Repeat until convergence:
  a. Calculate cells’ conditional log-likelihoods under each cluster using Algorithm 1 and the combined reference profiles matrix X = [X^(free)^, X^(ref)^].
  b. For each cohort of cells, compute posterior log-likelihoods using Algorithm 2.
  c. Update the cell type assignments by assigning each cell to the cell type under which it has the highest posterior log-likelihood.
  d. Update the profiles matrix X^(free)^ using Algorithm 3. Do not update the fixed reference profiles matrix X^(ref)^.
3. Declare convergence when fewer than 1/10^4^ cells change cell type, with a change in cell type defined as 1. A different cell type has the highest posterior probability, and 2. The change in posterior probability for the new cell type is >0.05.

### Subsetting by sketching

To help capture rare cell types, Insitutype modifies geometric sketching (2019), a biased subsampling technique that over-represents cells with rare expression patterns. Our algorithm for fast subsampling that favors rare cell types is below:

#### Algorithm 6

1. Take as input:
  1. The PCA reduction of the observed expression matrix
  2. The number of cells to sample
  3. The fractional difference between the total number of cells and the desired number of bins
  4. The maximum number of iterations used to find the desired number of bins
2. Scale each PC between 0 and 1 using the minimum and maximum values.
3. Repeat until the number of bins that contain a cell surpasses the minimum number of bins
  1. Subdivide each scaled PC into n ranges where n is 1 + the current number of iterations.
  2. With bins represented by each unique combination of ranges across all PCs, find the number of bins that contain cells.
4. Sample cells with probability equal to the inverse of their bin size.

### Criterion for selecting the number of clusters

To compare solutions under two different cluster numbers, we use the Akaike Information Criterion (AIC). We calculate AIC as follows:

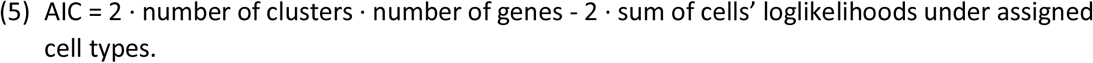

The first term captures the number of parameters, and the second term captures the total log-likelihood.

Alternatively, the Bayesian Information Criterion can be used:

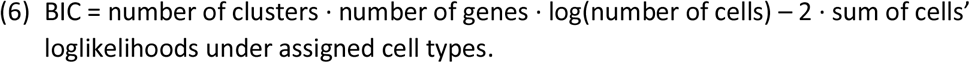

To speed computation, we evaluate cluster numbers using a subset of 10,000 cells. For each cluster number under consideration, we perform Insitutype on the 10,000-cell subset, then record the resulting AIC. The cluster number returning the lowest AIC is taken.

### Insitutype algorithm for unsupervised and semi-supervised clustering

To enable scaling to very high numbers of cells, the Insitutype algorithm coordinates calls to Algorithm 5 over subsets of the data.

#### Algorithm 7

1. Take as input:
  a. The observed expression matrix Y
  b. An initial matrix of expected cell type expression profiles X (potentially carefully pre-specified, or potentially derived from random cluster assignments.)
  c. Estimated per-cell background, usually estimated as described in the section, “Estimating individual cell’s background rates”.
  d. Mean negative control counts per cell.
  e. Optionally, a matrix of reference profiles.
  f. A vector specifying cells’ “cohort” membership, potentially all the same.
  g. The number of clusters, or a range of cluster numbers to consider.
2. Try multiple random starting points in small subsamples.
  a. Select 5 subsamples of 10,000 cells using Algorithm 6.
  b. Use Algorithm 5 to cluster each subsample, and save the profiles matrix X from each solution.
  c. In a new subsample of 10,000 cells, evaluate the log-likelihood of each cell under the profile matrix X from each clustering solution. Retain the profile matrix that produces the highest total log-likelihood.
3. Refine the clustering solution in a larger subsample.
  a. Take a subsample of 20,000 cells using Algorithm 6.
  b. Run Algorithm 5, initialized with the profile matrix from step 2.
4. Further refine the clustering solution in a very large subsample.
  a. Take a subsample of 100,000 cells using Algorithm 6.
  b. Run Algorithm 5, initialized with the profile matrix from step 3.
5. Use the profiles matrix from step 4 to perform supervised cell typing of all cells, using Algorithm 4.

### Updating reference profiles to account for platform effects

Due to platform effects and between-tissue differences, reference profile matrices from scRNA-seq differ from the cell type profiles as measured by a spatial platform, effectively adding noise to the reference profiles. This noise has not proved problematic for supervised cell typing. However, during semi-supervised clustering, noisy reference profiles cannot fit the data as well as free profiles derived during clustering, and the free clusters steal cells from the reference cell types.

To avoid bias against reference cell types during semi-supervised clustering, the reference profile matrix must be updated to better fit the spatial data. This update is performed as follows:

#### Algorithm 8

1. Identify “anchor cells”: cells that can confidently be assigned to a reference cell type. Anchor cells can be hand-selected based on marker genes or morphology, or they can be selected in a data-driven approach using Algorithm 9.
2. Estimate the mean profile of each cell type’s anchors using Algorithm 3.

### Choosing anchor cells

Anchor cells can be automatically nominated using the following algorithm:

#### Algorithm 9

1. Compute each cell’s log-likelihood under each reference profile using Algorithm 1.
2. Compute each cell’s cosine similarity to each reference profile.
3. For each cell type, take as anchors all cells with cosine similarity > 0.3 and with log-likelihood ratio to other cell types > 0.01 · total transcripts.

It is often useful to use cells’ immunofluorescence values as an additional filter.

### Analysis of cell pellet array to evaluate impact of orthogonal data types

Each cell’s image data was summarized using the mean intensity of the immunofluorescence markers for PanCK, CD45, CD3, and CD298. When scoring error rates, clusters were assigned to whichever cell pellet was most abundant within them.

### Benchmarking analysis

The cell pellet array dataset used for benchmarking analysis consisted of 52,518 cells from 13 cell lines, and the assay profiled 960 genes. The first downsampled dataset contains a random sample of 50% of the genes. The second downsampled dataset contains the 10% most highly variable genes, as defined by the getTopHVGs function from the R scran package.

For unsupervised clustering, Institutype was compared against K-means, Mclust, Leiden, and SC3. As the data contains 13 distinct cell lines, all clustering methods were run with 13 clusters. The metric used is the adjusted Rand index (ARI), where a high ARI indicates high similarity between the true cell type labels and the clustering partition. ARI equals 1 when there is perfect correspondence and takes a value near 0 for a random partition. For K-means, Mclust, and Leiden, principal components analysis was first performed, and the top 50 principal components were used in clustering analysis. K-means was run using default settings from the kmeans function in the R stats package. Mclust was run using the Mclust function with the “EEE” model parameter. Leiden was run using the cluster_leiden function from the igraph package on a shared nearest-neighbor graph with 20 neighbors. The resolution parameter was tuned to generate 13 clusters. SC3 was run using the recommended workflow for large datasets, which involves running the sc3 function on a random subset of 5,000 cells to train a model that can then be used to predict the remaining cells via the sc3_run_svm function.

For supervised cell typing, Insitutype was compared against SingleR, CHETAH, and Seurat label transfer. The metric used is error rate, calculated by comparing the inferred and true cell type label for each cell. The reference dataset was another cell pellet array dataset with 36,300 new cells from the same 13 cell lines. SingleR was run using default parameters. Seurat label transfer was run using the recommended workflow, including running the NormalizeData function on both the reference and test datasets, the FindTransferAnchors function, and then TransferData function. CHETAH was run using the recommended workflow. The reference dataset was normalized using the logNormCounts function and t-SNE was performed on the test dataset using the runTSNE function. Both functions are available in the R scater package.

### Analysis of lupus nephritis sample

Each cell’s image data was summarized using metrics collected by the default CosMx pipeline: area, aspect ratio, and mean intensity of the immunofluorescence markers for PanCK, CD45, CD3, CD298, and DAPI. Each cell’s spatial context was defined by taking as the mean expression profile of its 50 closest neighbors; these mean expression profiles were then reduced to their first 10 principal components. Insitutype’s default algorithm then defined cohorts based on these 17 variables. Reference profiles derived from the Human Cell Atlas adult kidney dataset (Regev 2017) were accessed from NanoString’s cell profile library (Danaher 2022, https://github.com/Nanostring-Biostats/CellProfileLibrary).

### Processing of non-small cell lung cancer data

The NSCLC CosMx dataset was downloaded from the NanoString website (https://nanostring.com/products/cosmx-spatial-molecular-imager/ffpe-dataset/). Cohorting was performed by applying Insitutype’s default cohorting algorithm to the following cell image attributes: area, aspect ratio, and mean intensities from the immunofluorescence stains for CD298, PanCK, DAPI, CD45, and CD3.

### Preparation of cell pellet array

Custom FFPE cell pellet arrays of 16 cell lines for RNA assay were made with A-FLX™ FFPE CELL PELLET by Acepix Bioscience, Inc. All cell lines were originally sourced from ATCC. The 16 cell lines for RNA assay include CCRF-CEM, COLO205, DU145, EKVX, HCT116, HL60, HOP92, HS578T, IGROV1, M14, MDA-MB-468, MOLT4, PC3, RPMI-8226, SKMEL2, SUDHL4.

Data collection failed for the FOVs from the cell lines HL60, IGROV1, and MOLT4, and so these cell lines were omitted from all analyses.

### Preparation of lupus nephritis sample

Slides were baked overnight at 60°C to ensure sample adherence to the glass slides. Then the baked samples went through deparaffinization, proteinase K digestion, and heat-induced epitope retrieval (HEIR) procedures to expose target RNAs and epitopes using the Leica Bond Rx system. 3 μg/ml of proteinase K incubation at 40°C for 30 min and HEIR at 100°C for 15 min in Leica buffer ER1 conditions were used. The samples were rinsed with DEPC H_2_O twice before incubating in 1:1000 diluted fiducials (Bangs Laboratory) in 2X SSCT solution for 5 min at room temperature. Excessive fiducials were removed by rinsing the samples with 1X PBS, followed by fixation with 10% neutral buffered formalin (NBF) for 5 min at room temperature. Fixed samples were rinsed with Tris-glycine buffer (0.1M glycine, 0.1M Tris-base in DEPC H_2_O) and 1X PBS for 5 min each before blocked using 100 mM N-succinimidyl acetate (NHS-acetate, ThermoFisher) in NHS-acetate buffer (0.1M NaP+0.1% Tween PH8 in DEPC H_2_O) for 15 min at room temperature. Prepared samples were rinsed with 2X saline sodium citrate (SSC) for 5 min and then Adhesive SecureSeal Hybridization Chamber (Grace Bio-Labs) was placed to cover the samples.

980-plex RNA ISH probes were denatured at 95°C for 2 min and then placed on ice before making ISH probe mix (1 nM ISH probes, 1X Buffer R, 0.1 U/μL SUPERaseIn™ in DEPC H_2_O). The ISH probe mix was pipetted into the hybridization chamber and the chamber was sealed using adhesive tape. Hybridization occurred at 37°C overnight after sealing the chamber to prevent evaporation. After the overnight hybridization, samples were washed with 50% formamide (VWR) in 2X SSC at 37°C for 25 min for 2 times, rinsed with 2XSSC for 2 min for 2 times at room temperature, and then blocked with 100 mM NHS-acetate for 15 min. After blocking, washed the samples twice using 2X SSC for 2 min at room temperature. Custom-made slide cover was attached to the sample slide to form a flow cell.

Target RNA readout on the CosMx instrument was performed following published protocols (He 2022). In brief, the assembled flow cell was loaded onto the CosMx instrument and the TMA samples were washed with reporter wash buffer to rinse the samples and remove air bubbles. Once thes flow cell was loaded onto the instrument correctly, the entire flow cell was scanned (CosMx preview scan) and 15-25 fields of view (FOVs) were placed on the TMA cores for RNA readout. RNA readout cycle was started by flowing 100 ul of Reporter Pool 1 into the flow cell and incubated for 15 min. After incubation, 1 mL of Reporter Wash Buffer was flowed into the flow cell to wash out the unbound reporter probes, followed by replacing Reporter Wash buffer with Imaging buffer prior to imaging. Nine Z-stack images (0.8 um step size) of each FOV were acquired and then fluorophores on the reporter were UV cleaved and washed off with Strip Wash buffer. This fluidic and imaging procedure was repeated for the 16 reporter pools, and the 16-round of reporter hybridization-imaging cycle was repeated 9 times to increase RNA detection sensitivity.

After 9 cycles of RNA readout, TMA samples were incubated with 4 fluorophore-conjugated antibody cocktail against CD298, PanCK, CD45, and CD3 proteins and DAPI in the same instrument for 1 hr. Nine Z-stack images for 5 channels (4 antibodies and DAPI) were captured after unbound antibodies and DAPI were washed with Reporter washing buffer and then the flow cell was filled with Imaging buffer.

Raw image processing and feature extraction were performed using in-house data processing pipeline (He 2022). The pipeline consists of 3 main steps – registration, feature detection, and localization. For the first step, 3D rigid image registration was made using fiducials embedded in the TMA samples and then matched with the fixed image reference established at the beginning of CosMx run to correct any shift happening during the RNA readouts. Secondly, the RNA image analysis algorithm found reporter signature locations in X, Y, and Z axes along with the assigned confidence and then recorded all features into a single list. Lastly, individual target transcript XYZ location information was extracted and written into a table by secondary analysis algorithm.

### Cell segmentation

Immunostaining with DAPI Z-stack images was used for drawing cell boundary on the samples (He 2022). A cell segmentation pipeline (Stringer 2021) was used to assign transcripts to cells. The segmentation algorithm also maps individual transcripts to the subcellular compartments such as nuclei, cytoplasm, and membrane. Additional outputs of the algorithm include cell area, aspect ratio, and transcript statistics per cell. The transcript profile of individual cells was generated by combining target transcript location and cell segmentation information, which was fed into tertiary data analysis.

## Supplementation information

### Biased subsampling enhances detection of rare cell types

Detection of rare cell types is a persistent challenge in single cell transcriptomics analyses (Kiselev 2019). An opportunity is presented by the small subsets in Insitutype’s initial iterations. If these small subsets can be biased towards rare cell types, the early iterations will more easily fit those cell types, establishing them as clusters that will persist through the final iterations.

To enrich rare cell types within subsamples, Insitutype employs sketching (Hie 2019), a class of rough but very efficient clustering algorithms. It partitions the data into thousands of clusters, then samples equally from each cluster.

The sketching algorithm is demonstrated in Supplemental Fig. 1. In a kidney core biopsy, 61073 cells were assigned to 33 cell types by comparison to reference profiles from the Human Cell Atlas (Regev 2017). Insitutype’s sketching algorithm was then used to select a subset of 2000 cells. Rare cell types were enriched in the sketching-derived subsample: the Gini coefficient of the 33 granular cell types dropped from 0.70 in the complete data to 0.62 in the subsampled data, and the Gini coefficient of 10 cell type supersets dropped from 0.65 to 0.47. The least abundant cell superset, lymphoid lineage cells, rose from just 0.62% (Wilson 95% confidence interval 0.55% – 0.68%) of the complete dataset to 2.60% (1.99%– 3.93%) in the subsample.

**Supplementary Figure 1:**
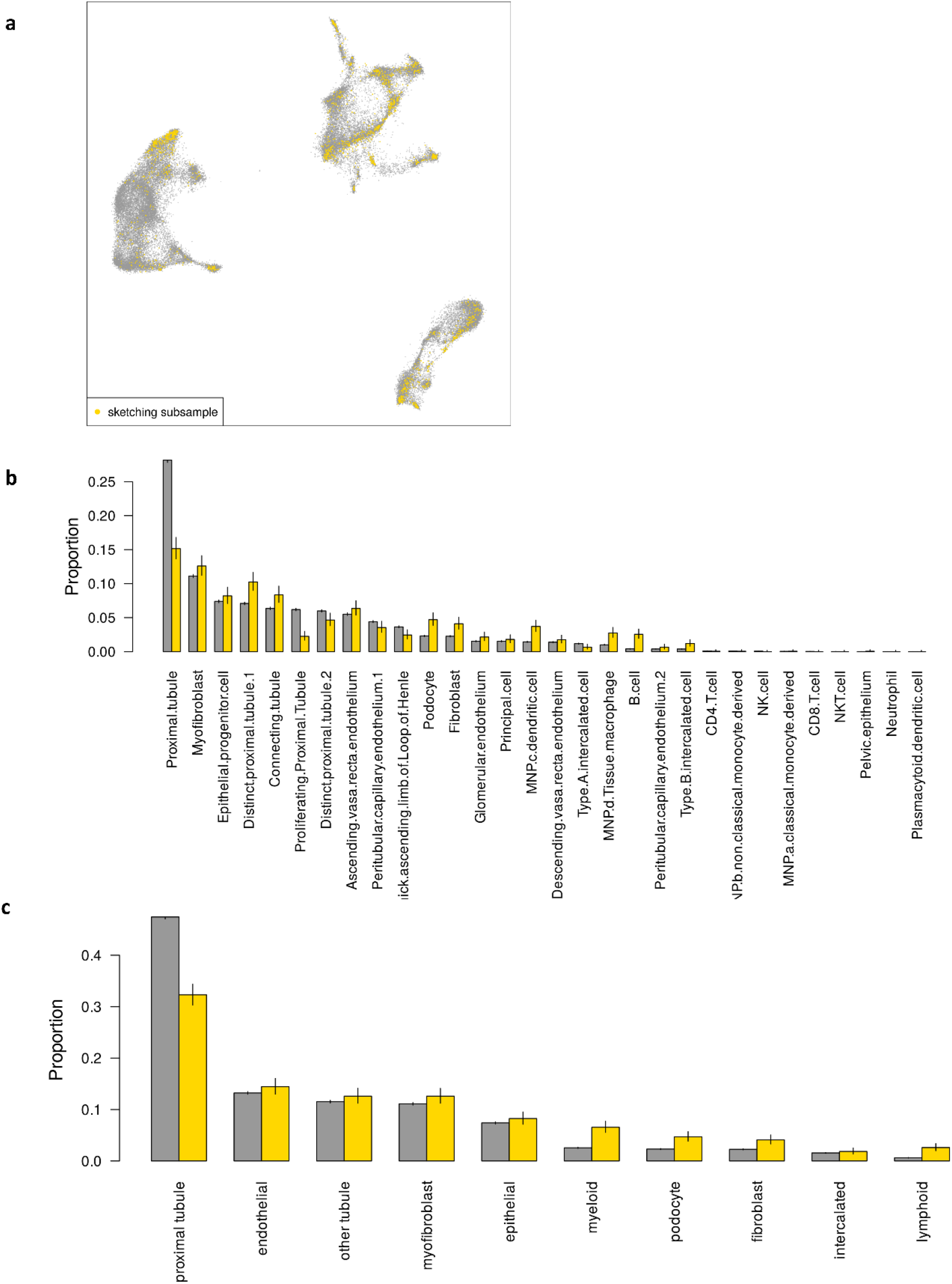
sketching to overrepresent rare cell types. **a**. UMAP of 61073 kidney cells. Black points show the selected subset of 2000 cells. **b**. Frequency of 33 cell types in the full dataset and in the sketching-derived subset. **c**. Frequency of 10 cell type supersets in the full dataset and in the sketching-derived subset.

**Supplemental Figure 2.**
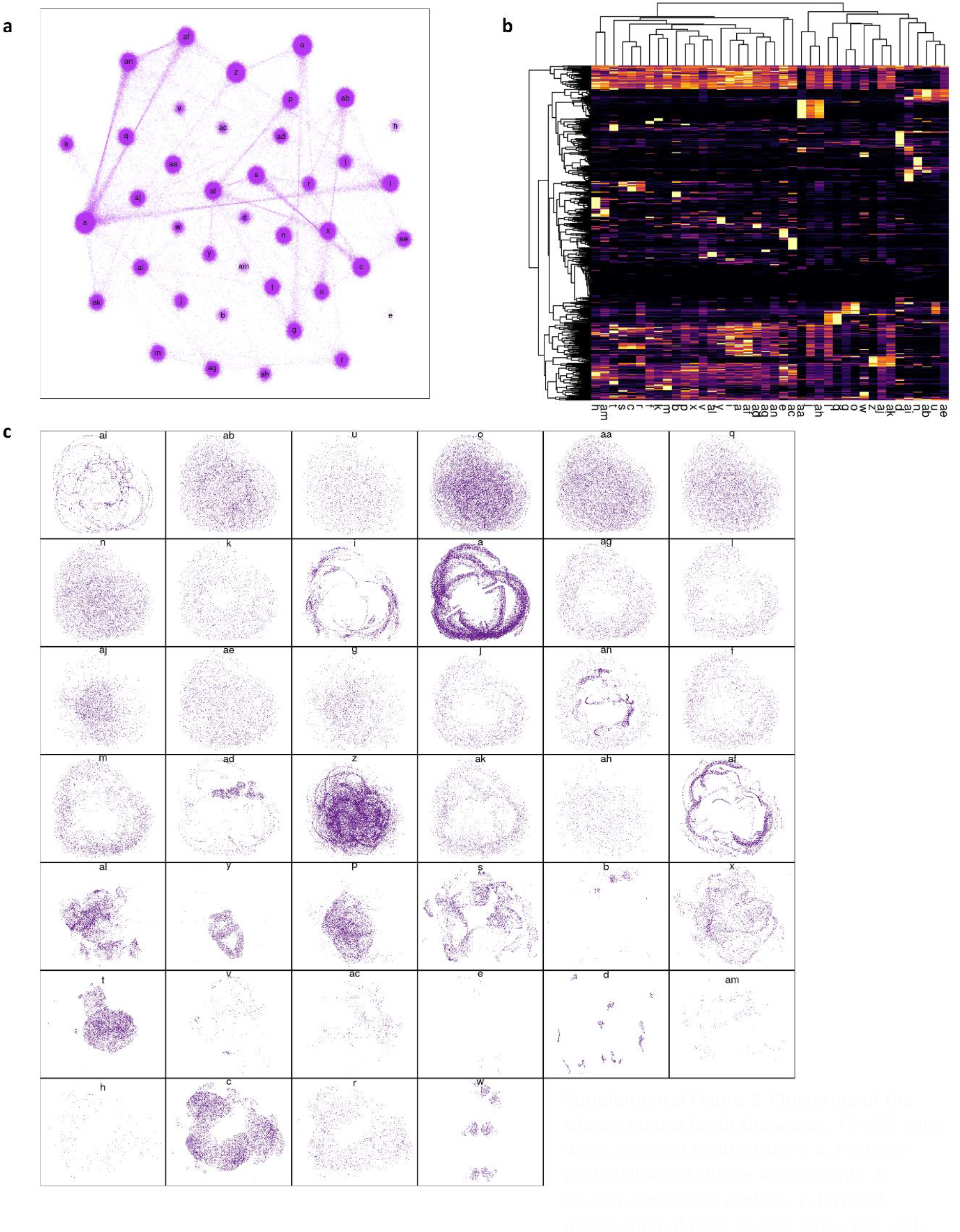
Clustering of the Vizgen mouse brain showcase. 734696 cells were assigned to 40 clusters. **a**. Posterior probabilities of cluster assignments. **b**. Cluster expression profiles. **c**. Physical distribution of each cluster. Only cells with posterior probability= 1 are shown.

**Supplemental Figure 3:**
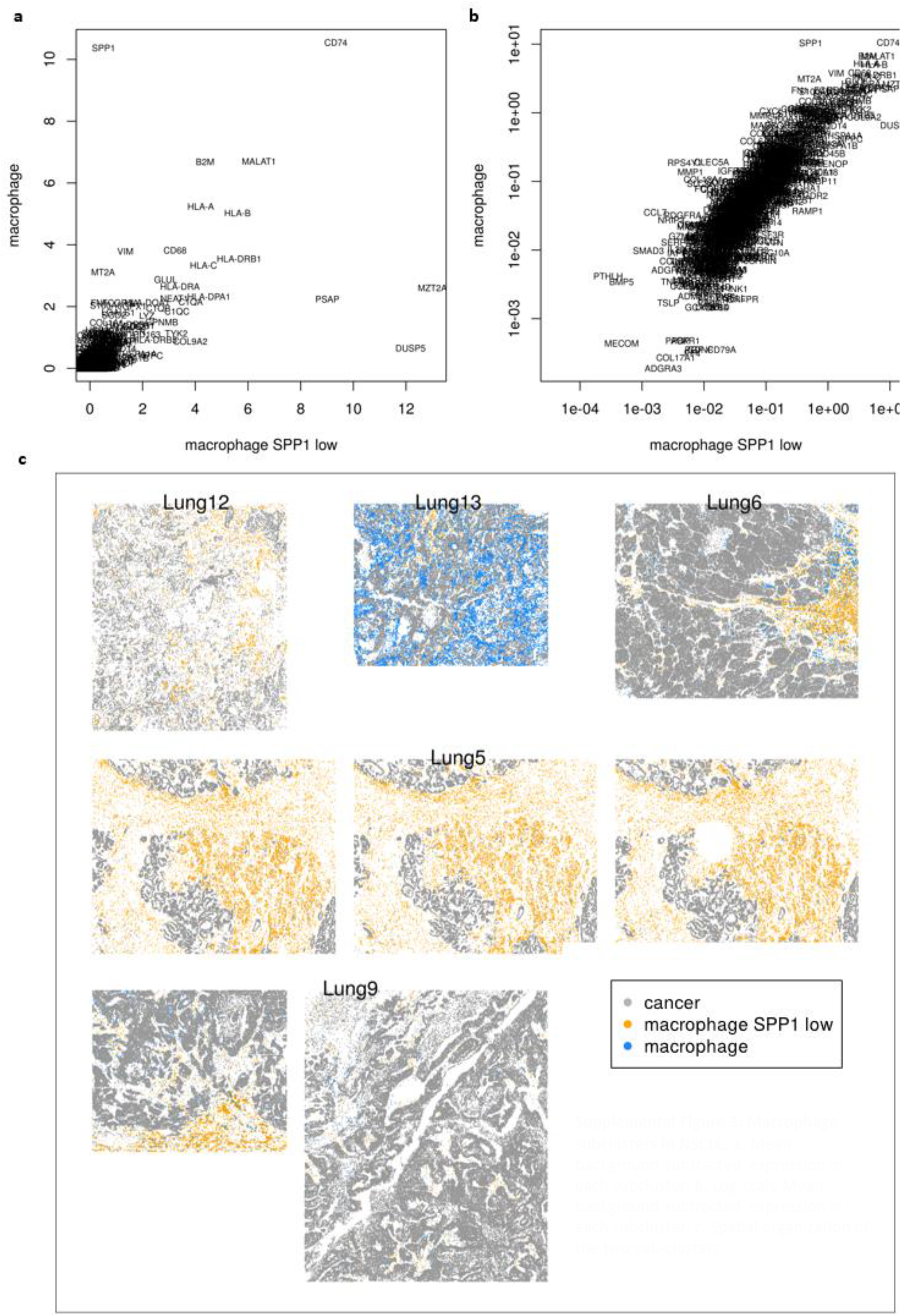
Macrophage subclusters in NSCLC. **a**. Mean background-subtracted expression in each subcluster. **b**. Log-scale Mean background-subtracted expression in each subcluster. **c**. Spatial organization of the two sub-clusters.

**Supplemental Figure 4:**
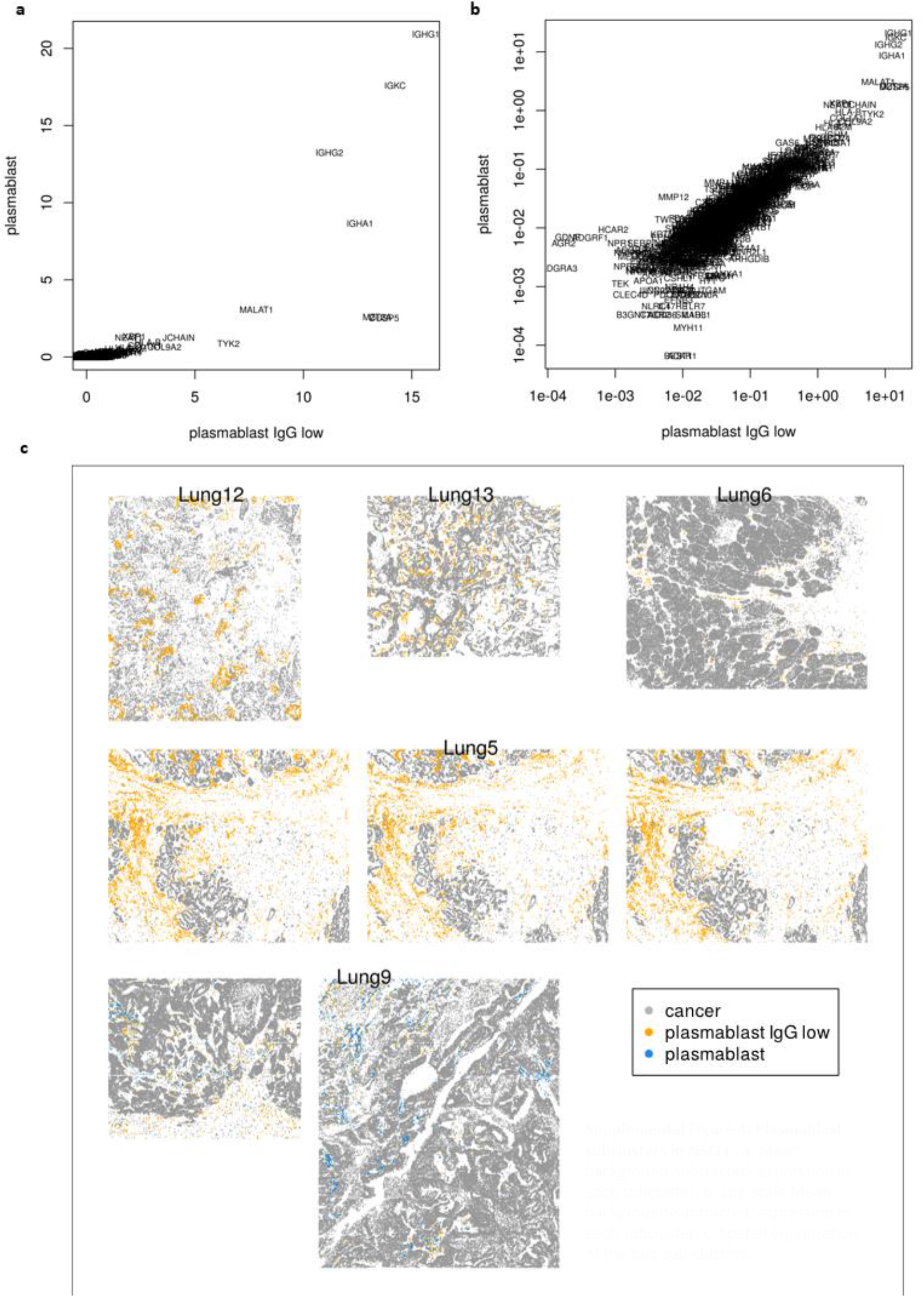
Plasmablast subclusters in NSCLC. **a**. Mean background-subtracted expression in each subcluster. **b**. Log-scale Mean background-subtracted expression in each subcluster. **c**. Spatial organization of the two sub-clusters.

**Supplemental Figure 5:**
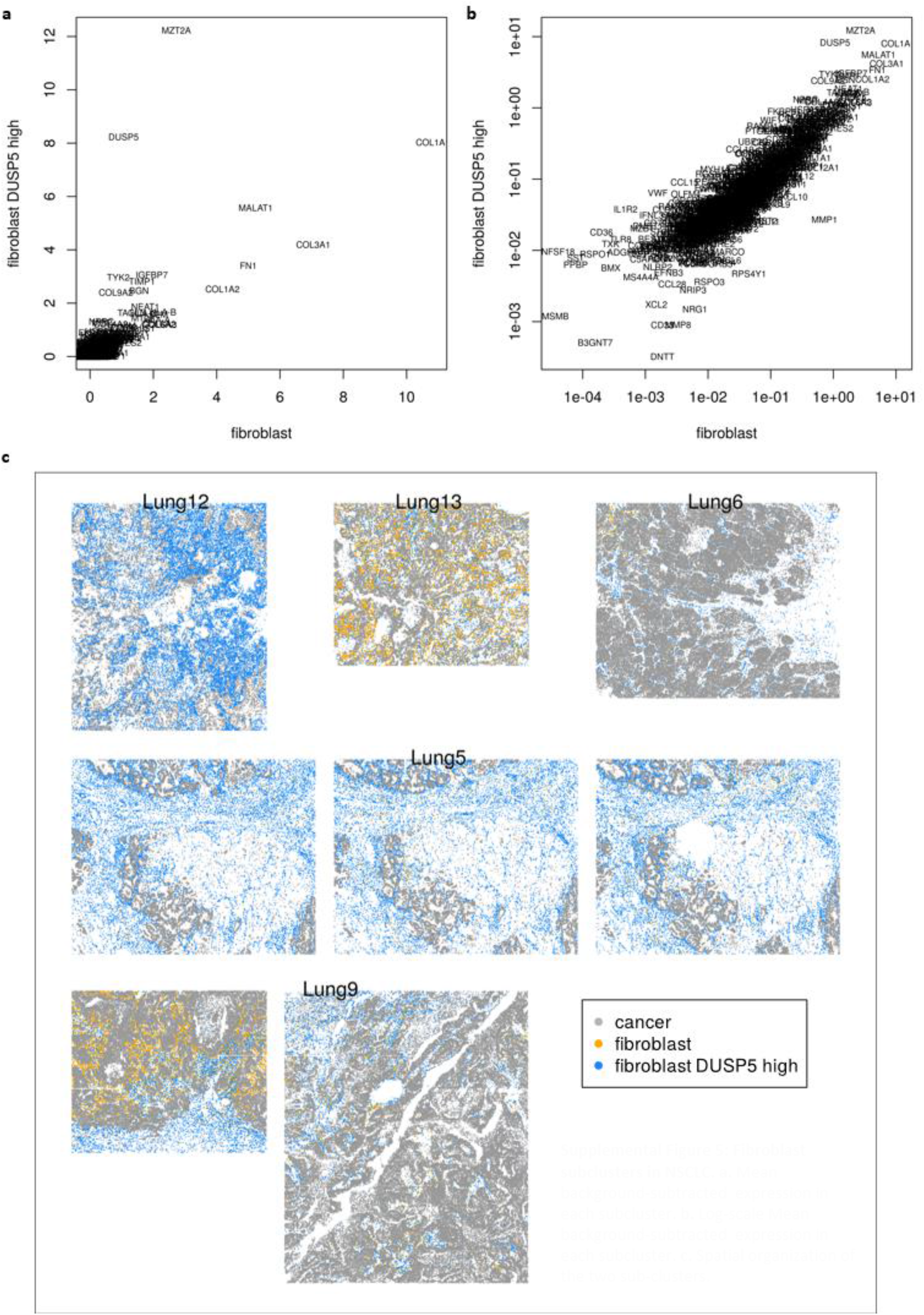
Fibroblast subclusters in NSCLC. **a**. Mean background-subtracted expression in each subcluster. **b**. Log-scale Mean background-subtracted expression in each subcluster. **c**. Spatial organization of the two sub-clusters.

**Supplemental Figure 6:**
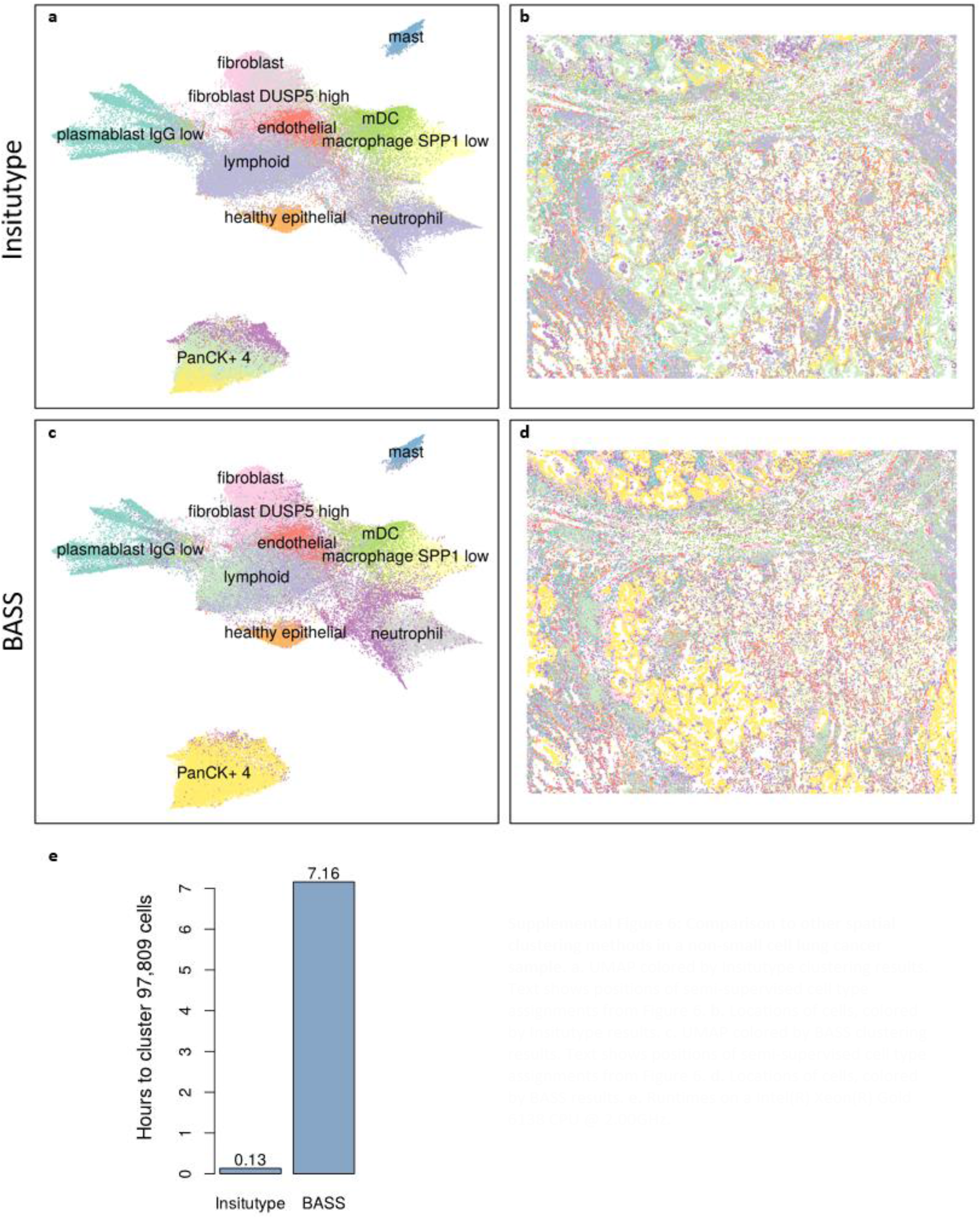
Comparison to other spatial clustering methods in a non-small cell lung cancer sample. **a**. UMAP colored by lnsitutype clustering results. Text shows positions of semi-supervised cell type assignments from Figure 6. **b**. Locations of cells, colored by Insitutype results. **c**. UMAP colored by BASS clustering results. Text shows positions of semi-supervised cell type assignments from Figure 6. **d**. Locations of cells, colored by BASS results. **e**. Runtimes on a lntel(R) Xeon(R) Gold 6138 CPU @ 2.00GHz.

